# Difference in accumulation pattern of the allergens within the same PR10 family in transgenic rice

**DOI:** 10.1101/356972

**Authors:** Fumio Takaiwa, Yuko Ogo, Yuhya Wakasa

## Abstract

Apple food allergen Mal d 1 and birch pollen allergen Bet v 1 belong to the same pathogen related protein 10 (PR10) family. When each of these allergens was expressed as a secretory protein by fusion with the GFP reporter in transgenic rice by ligating an N terminal signal peptide and a C terminal KDEL ER retention signal under the control of the maize ubiquitin constitutive promoter, the GFP:Mald1 highly accumulated in various tissues, whereas the accumulation level of GFP:Betv1 was remarkably reduced in vegetative tissues except for seed. Analysis by RT-PCR exhibited that there was little difference in transcript levels between them, indicating the involvement of post-transcriptional regulation. To investigate the cause of such difference in accumulation levels, deletion analysis of the Mal d 1 and domain swapping between them were carried out in transgenic rice. These results showed that the region between positions 41-90 in the Mal d 1 is predominantly implicated in higher level accumulation in vegetative tissues as well as seed compared with the Bet v 1. It is notable that GFP:Mald1 directed by the ubiquitin promoter is deposited in huge PBs in aleurone layer rather than starchy endosperm.

**Highlight:** Specific region of PR10 proteins is mainly implicated in their stability in vegetative tissues when expressed in transgenic rice.

## Introduction

Mal d 1 is the major allergen of apple fruit eliciting oral allergy syndrome (OAS) such as itching and swelling of lips, tongue and throat, which belongs to the family of pathogen-related 10 (PR10) proteins within the Bet v 1 superfamily (Mari *et al.* 2005; Fernandes *et al*, 2013). It has been reported that more than 70% of birch pollen-allergenic patients sensitizing to the major birch pollen allergen Bet v 1 display OAS after apple consumption (Geroldinger-Simic *et al.*, 2011). Namely, the OAS by apple is provoked by the allergic cross-reaction between the Mal d 1 and Bet v 1 specific IgE, indicating the presence of common sequences (IgE epitopes) and tertiary structure required for its binding to the Bet v 1 specific IgE. (Ebner *et al.*, 1991; Geroldinger-Simic *et al.*, 2011).

Mal d 1 is composed of 159 amino acids with the estimated molecular mass of 17.5 kDa. Its subcellular localization is estimated to be intracellular and cytoplasmic like the other PR10 proteins, since there is no signal peptide sequence directing to the endomembrane system as a secretory protein (Fernandes *et al.* 2013). Mal d 1 is mainly present in the pulp and peel of apple, but is not only expressed in apple fruit but also in various vegetative tissues and seed (Beuning *et al.*, 2004; Marzban *et al.*, 2005). Although the biological function of Mal d 1 remains unknown well, its expression has been studied to respond to fungal or bacterial infection and also to be activated by stress and storage, suggesting that it participates in defense mechanism against pathogens or abiotic stress like other PR10 proteins (Fernandes *et al.* 2013).

Mal d 1 is encoded by a multigene family composed of 31 members, which are divided into four groups based on amino acid sequence similarities, Mal d 1.01, Mald d 1.02, Mal d 1.03 and Mal d 1.04 (Mal d 1 a-d) (Pagliarani *et al.*, 2013). Mal d 1.01 and Mal d 1.02 are predominantly expressed in apple fruit (peel and pulp) (Marzban *et al.*, 2005). Amino acid sequence identities between Mal d 1 isoforms and Bet v 1 range from 55 to 68 % (Ma *et al.*, 2006).

Rice seed provides good production platform for high-value recombinant proteins in terms of scalability, safety, stability and cost-effectiveness (Stoger *et al.*, 2005; Takaiwa *et al.*, 2015). Especially, when recombinant proteins are produced as secretory protein and then deposited into protein-bodies (PBs) in the endosperm cells, they are highly and stably accumulated (Takaiwa *et al.*, 2017). Moreover, when orally administered, it has been known that recombinant proteins bio-encapsulated in PBs are effectively delivered to small intestine without degradation with digestive enzymes in gastrointestinal tract, providing an ideal delivery system to gut-associated lymphoid tissues (GALT) as oral mucosal vaccines (Hofbauer and Stoger, 2013; Takaiwa *et al.*, 2015).

We previously generated transgenic rice seeds containing versatile hypo-allergenic derivatives of major birch pollen allergen Bet v 1 referred to as tree pollen chimera7 (TPC7) or TPC9, which have been developed as allergy vaccines (tolerogens) against the major allergens of trees belonging to the Fagales order Bet v 1 family for allergen-specific immunotherapy (Wang *et al.*, 2013). When these native Bet v 1 and its derivatives (TPC7 and TPC9) were expressed as secretory protein in rice seeds under the control of the endosperm-specific glutelin *GluB-1* promoter by ligating the glutelin GluB-1 signal peptide and the KDEL ER retention signal to their N- and C-termini, they were deposited in remarkably huge spherical ER-derived PBs designated as TPC7 bodies with the size of >10 μm (Wang *et al.*, 2013; Ogo *et al.*, 2014). Furthermore, their accumulation levels reached about 23% of total seed proteins (Ogo *et al.*, 2014). Such ability to form huge PBs is expected to have a potential to produce high amounts of foreign recombinant proteins by fusion to them. Moreover, it is interesting to examine whether properties yielding such huge PB formation and high productivity are retained in vegetative tissues, when expressed in vegetative tissues.

In this study, we examined whether Mal d 1, Bet v 1 and TPC7 within the same PR10 family are deposited in huge ER-derived PBs in various tissues of transgenic rice. When expressed under the control of the maize ubiquitin constitutive promoter. it was shown that the Mal d 1 stably and highly accumulates in vegetative tissues as well as seed, whereas the accumulation levels of the Bet v 1 and TPC7 are very low in vegetative tissues except for seed. Analysis by deletion and domain-swapping between the Mal d 1 and Bet v 1 demonstrated that this difference is mainly attributed to the region between positions 41-90.

## Methods

### Construction of expression vectors and generation of transgenic rice

The DNA sequences encoding Mal d 1, Bet v 1 and TPC7 were optimized for translation based on the codons frequently used in rice seed storage protein gens and their DNA sequences were synthesized by GenScript Corporation (NJ, USA). The N terminal signal peptide of the glutelin GluB-1 that is required for translocation into the ER and the KDEL ER retention signals were attached to their N and C termini, respectively. The gene encoding the green fluorescence protein (GFP) reporter was inserted in frame between the GluB-1 signal and these PR10 related genes. In order to express specifically in the endosperm or constitutively in vegetative tissues, 2.3kb glutelin *GluB-1* promoter or maize ubiquitin promoter were ligated upstream of these three fusion constructs, which were followed by the 0.65kb of glutelin *GluB-1* terminator (Fig. 1B).

**Figure 1.**
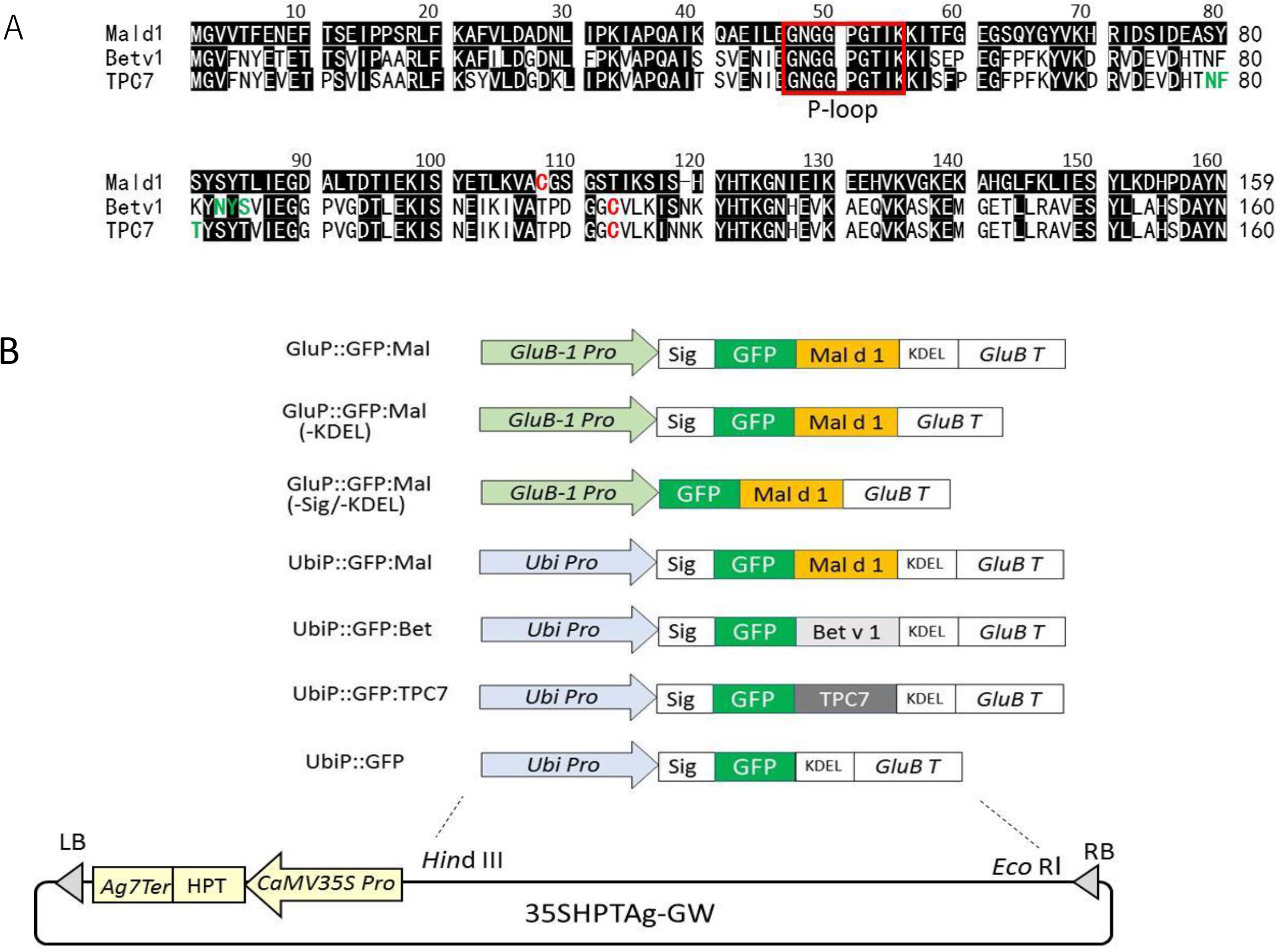
Gene constructs used for expression of Mal d 1, Bet v 1 and TPC7 in transgenic rice. **(A)** Alignment of Mal d 1, Bet v 1 and TPC7 amino acid sequences. Cysteine residue and glycosylation sites are indicated by red and green colors. Red square represents P loop motif. **(B)** Schematic representation of the gene constructs used for expression of GFP, GFP:Mald1, GFP:Betv1 and GFP:TPC7 in transgenic rice. Mal d 1, Bet v 1 and TPC7 were expressed as fusion proteins with GFP under the control of the 2.3 kb glutelin *GluB-1* promoter and maize ubiquitin promoter. SP and *GluB-1* T represent the glutelin B1 signal peptide sequence and 0.65 kb 3’ untranslated region of *GluB-1*, respectively. KDEL, endoplasmic reticulum retention signal; HPT, hygromysin phosphotransferase gene; CaMV35S P, cauliflower mosaic virus 35S promoter; UbiP, maize ubiquitin gene promoter; RB, right border; LB, left border

These gene expression cassettes inserted into entry vector were transferred into
destination binary vector (p35SHPTAg7-GW) harboring the CaMV35S promoter::hygromycine phosphotransferase gene (*HPT*) as selectable marker gene via the MultiSite Gateway LR Clonase reaction (Invitrogen) (Wakasa *et al.*, 2006). These binary vector plasmids containing various expression constructs were introduced into rice genome via *Agrobacterium*-mediated transformation (Goto *et al.*, 1999).

For each construct, more than 16 independent transgenic lines (16-30 lines) were generated. Transgenic rice plants were selected by hygromycin resistance and grown in the closed control greenhouse (33°C, 16h light/8h dark cycle).

### SDS-PAGE and immunoblot analysis

Individual mature grain (four grains per independent transgenic line) and leaves and roots frozen in liquid nitrogen were ground into a fine powder using a Multibeads shocker (Yasui Kikai, Osaka, Japan). Total proteins were extracted from the powder of a mature grain, leaves and roots with 400 μl (grain) or 200 μl (leaves and roots) of urea-SDS buffer (50 mM Tris-HCl, pH 6.8, 8M urea, 4% SDS, 5% 2-mercaptoethanol (2-MER), 20% glycerol) as described previously (Tada *et al.*, 2003). After separation of total protein extract (2-5 μl) by electrophoresis on 13% SDS-PAGE, proteins were visualized by Coomassie Brilliant Blue (CBB)-R250 staining or were transferred to PVDF membranes (Millipore, Billerica, MA, USA) for immunodetection with rabbit anti-GFP or appropriate antibodies. Rabbit antibodies against TPC7, rice glutelin A (GluA), and rice chaperones of binding protein 1 (BiP1) and 4&5 (BiP4/5), protein disulfide isomerase-like 1-1 and 2-3 (PDIL1-1 and PDIL2-3), and calnexin (CNX) were prepared as described previously (Yasuda *et al.*, 2009; Oono *et al.*, 2010; Wakasa *et al.*, 2011; Wang *et al.*, 2013). For immunodetection, membranes were incubated with rabbit antibodies, followed by horseradish peroxidase-conjugated anti-rabbit secondary IgG antibodies (dilution 1:5000) (Cell Signaling Technology, Danvers, MA, USA). Bands were visualized using Clarity Western ECL detection kit (Bio-Rad).

### Confocal laser scanning microscopy and electron immunomicroscopy

Developing seeds were harvested 15-20 days after flowering (DAF) from various transgenic lines and then cross-sectioned with a DTK-1000 Microslicer (Dosaka EM) to approximately 150 μm in thickness for indirect immune-histochemical analysis (Yasuda *et al.* 2006). Mouse anti-GFP monoclonal antibody, rabbit anti-CNX, anti-PDIL1-1, anti-PDIL2/3, anti-BiP1-1, anti-BIP4/5, anti-TiP3 and anti-RM1 (14kDa prolamin) polyclonal antibodies were used to investigate their intracellular localization by Alexa-488 conjugated anti-mouse IgG antibody and Alexa-546 conjugated anti-rabbit IgG antibody. After immunostaining of samples, rhodamine B was used to stain ER-derived protein bodies (PB-Is). Samples cross-sections were spread on glass slides, sealed with a cover glass, and observed with a confocal laser scanning microscope (FLUOVEIW; OLYMPUS, Tokyo, Japan).

### RNA extraction and RT-PCR analysis

Total RNA was extracted from seeds as previously described (Takaiwa *et al.*, 1987). RT-PCR analysis was carried out using ReverTraAce qPCR RT Master Mix with gDNA Remover (TOYOBO, Osaka, Japan) with gene-specific primers for GFP and 17S rRNA.

### Extraction of recombinant proteins from mature seeds

Leaves and roots were collected from each transgenic rice plant and frozen instantly in liquid nitrogen and then powdered using a Multibeads shocker. Pulverized seed, leaves and roots were treated with ten volume of various extraction buffer (200 μl) for 20 min by vortex, and then centrifuged at 15000 rpm for 10 min. The supernatant was transferred to new tube and then equal volume of 2x sample buffer (125 mM Tris-HCl (pH6.8), 20% glycerol, 4%SDS) without 2-MER was added. Some aliquot (5ul) was applied and subjected to electrophoresis on 6 or 8 % SDS-PAGE.

## Results

### Endosperm-specific expression of Mal d 1 in transgenic rice

When GFP:Mald1 fusion protein were expressed as the secretory protein by ligating an N terminal glutelin signal peptide and a C terminal KDEL ER retention signal under the control of the endosperm-specific 2.3 kb glutelin *GluB-1* promoter, it accumulated at high levels in the endosperm of transgenic rice, which was detected as a visible band with the molecular mass of 50 kDa on CBB staining gel (Fig. S1). However, when the KDEL ER retention signal was removed, its accumulation (GFP:Mald1(-KDEL)) was significantly decreased. Furthermore, when produced in the cytoplasm by the absence of both the signal peptide and KDEL tag, its accumulation level (GFP:Mald1(-sig/-KDEL)) was remarkably reduced to undetectable level (Fig. S1). These results are in remarkable contrast with the expression of the inherent Mal d 1 in apple fruit, since it accumulates at relatively high level (1-20 μg/g FW) irrespective of absence of the signal peptide (Matthes and Schmitz-Eiberger, 2009).

On the other hand, when GFP:Mald1 was expressed as secretory protein in seeds, its effect on expression levels of various chaperones proteins was analyzed by immunoblotting using specific antibodies. In these transgenic seeds BiPs (BiP1 and BiP4/5) and PDIL2-3 were significantly increased compared with those in GFP:Mald1 (-sig/-KDEL) construct, indicating that ER stress was induced by production at high level as secretory protein (Fig. S1).

Subcellular localization of GFP:Mald1 with the presence or absence of the KDEL signal was investigated in maturing transgenic seeds by immuno-histological analysis using confocal laser scanning microscopy. As shown in Fig. 2A, the GFP:Mald1 containing the KDEL signal was deposited in a spherical huge ER-derived protein body called as TPC7 body (Wang *et al.*, 2013) with a diameter of 5-10 μm in maturing endosperm cells. By contrast, when the KDEL tag was removed, generation of giant PBs was lost and the GFP:Mald1(-KDEL) was sequestered in small granules in ER lumen (Fig. 2B).

**Figure 2.**
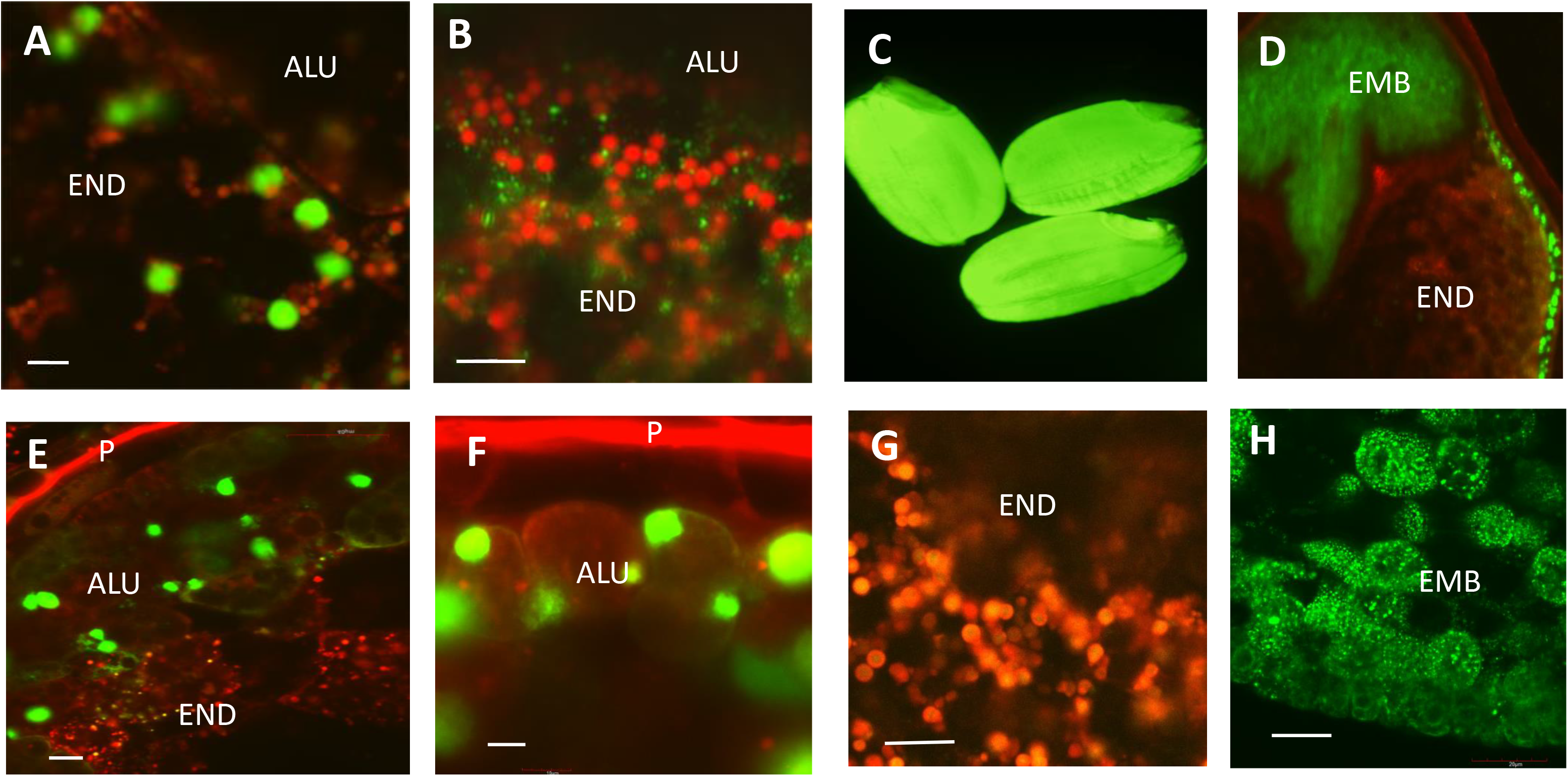
Expression pattern of GFP:Mald1 in transgenic rice seed. **(A and B)** Intracellular localization of GFP:Mald1 in transgenic rice seed harboring the GluP::GFP:Mal or GluP::GFP:Mal(-KDEL). **(C-H)** Expression pattern of GFP:Mald1 of transgenic rice seeds harboring UbiP::GFP:Mal and its intracellular localization in aleurone layers, starchy endosperm and embryo. ALU, aleurone layer; END, endosperm; EMB, embryo; p, pericalp thin Bars= 5 μm. thick Bars=100 μm

### Constitutive expression of Mal d 1, Bet v 1 and TPC7 in transgenic rice

The GFP:Mald1, GFP:Betv1 and GFP:TPC7 containing the GluB-1 signal peptide and KDEL ER retention signal at their N and C termini were expressed under the control of the maize ubiquitin promoter. The GFP:Mald1 accumulated in mature seeds at higher levels than the GFP:Betv1 and GFP:TPC7 (Fig. 3A). Next, individual whole seed expressed from these expression constructs was divided into the embryo and endosperm parts and their accumulation levels were examined by immunoblotting using anti-GFP antibody. As shown in Fig. 3C, accumulation levels of the GFP:Betv1 and GFP:TPC7 in most embryos were under detectable level, whereas the GFP:Mald1 highly accumulated in most embryos. These results are in contrast with that they accumulated at similar expression levels in their endosperm (Fig. 3B). Thus, difference in their accumulation levels in whole grains may be accounted for by their differences in embryos. On the other hand, when the GFP reporter containing the KDEL tag at its C terminus was directly expressed, it accumulated in both embryo and endosperm like the GFP:Mald1 construct (Fig. S2). These findings suggested that fusion to the Bet v 1 or TPC7 may give rise to unstability of their fusion products in embryos.

**Figure 3.**
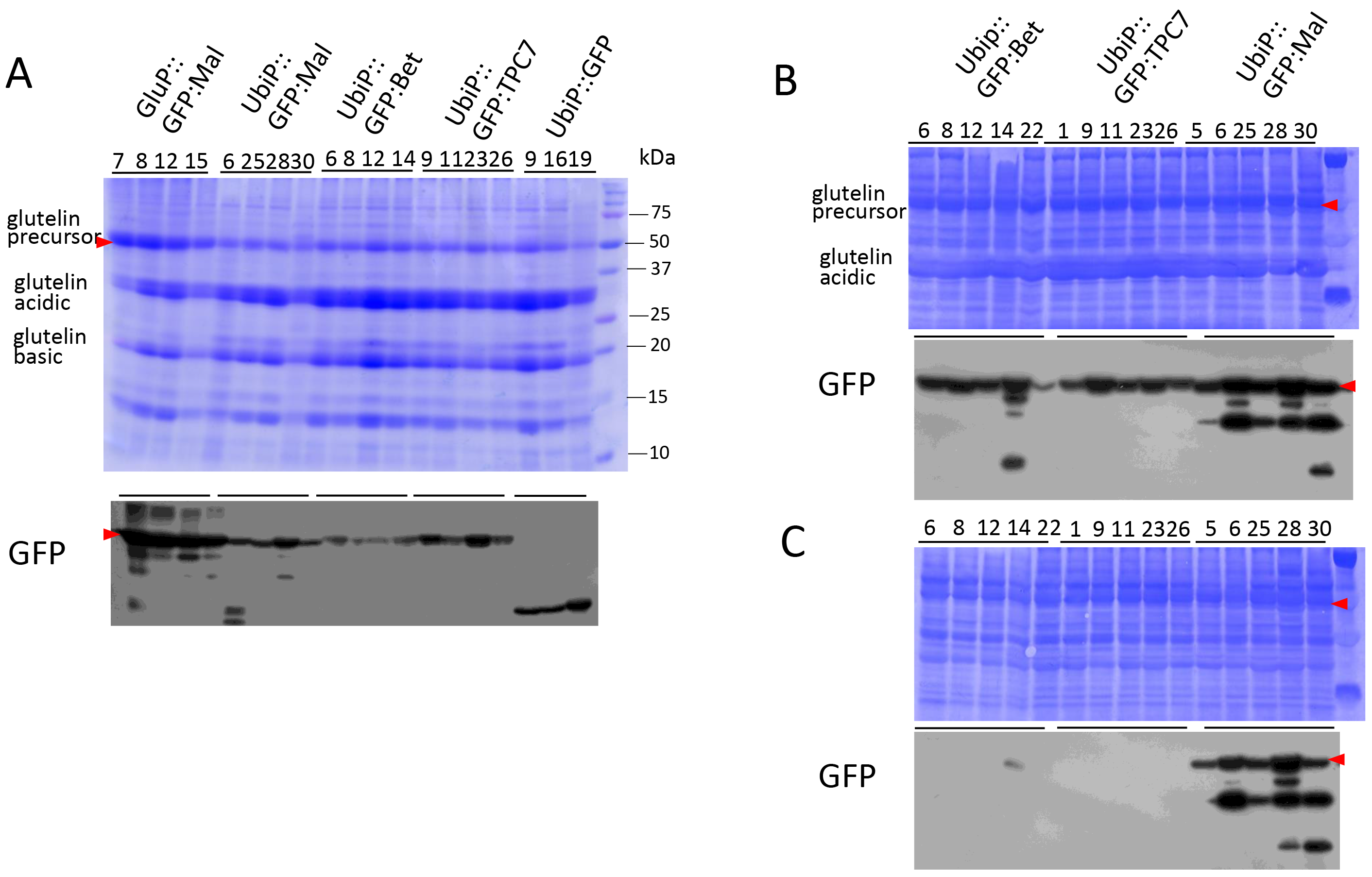
Immunoblot analysis of GFP:Mald1, GFP:Betv1 and GFP:TPC7 expressed in transgenic rice seeds under the control of the maize ubiquitin promoter. GFP:Mald1 was also expressed under the control of the glutelin *GluB-1* promoter for comparison. Total proteins were extracted from the whole seed **(A)**, endosperm **(B)** and embryo **(C)** with SDS-urea buffer, and subsequently separated by 13% SDS-PAGE. Accumulation levels were examined by immunoblotting with anti-GFP antibody. GFP fusion products with the predicted size of 50 kDa are indicated by arrow head. Degradation products of about 37 kDa and 20 kDa are detected in high expression lines.

When accumulation levels of the GFP:Mald1 in mature seeds that were directed by the endosperm-specific glutelin *GluB-1* promoter were compared with those that were driven by the ubiquitin promoter, the endosperm-specific promoter led to higher levels of its accumulation (Fig. 3A). This result may reflect the difference in promoter activities of maturing seeds between them (Qu and Takaiwa, 2004).

Next, accumulation levels of the GFP:Mald1, GFP:Betv1 and GFP:TPC7 in leaves and roots were examined by immunoblotting using anti-GFP antibody. It should be noted that these GFP fusion products with the size of 50kDa is processed (degraded) into 37-39 kDa in vegetative tissues, when analyzed by immunoblotting using anti-GFP antibody. As shown in Fig. 4A, GFP:Mald1 highly accumulated in leaves and roots, whereas the accumulation levels of GFP:Betv1 and GFP:TPC7 were very low. Especially, their accumulation levels in roots were much lower than those of GFP:Mald1. These results suggested that the GFP:Betv1 and GFP:TPC7 products are unstable in vegetative tissues in a similar manner to their expression in their embryos. Therefore, the Mal d 1 may act as stabilizing sequence by protecting fusion proteins from degradation or the Bet v 1 and TPC7 may cause unstability in vegetative tissues.

**Figure 4.**
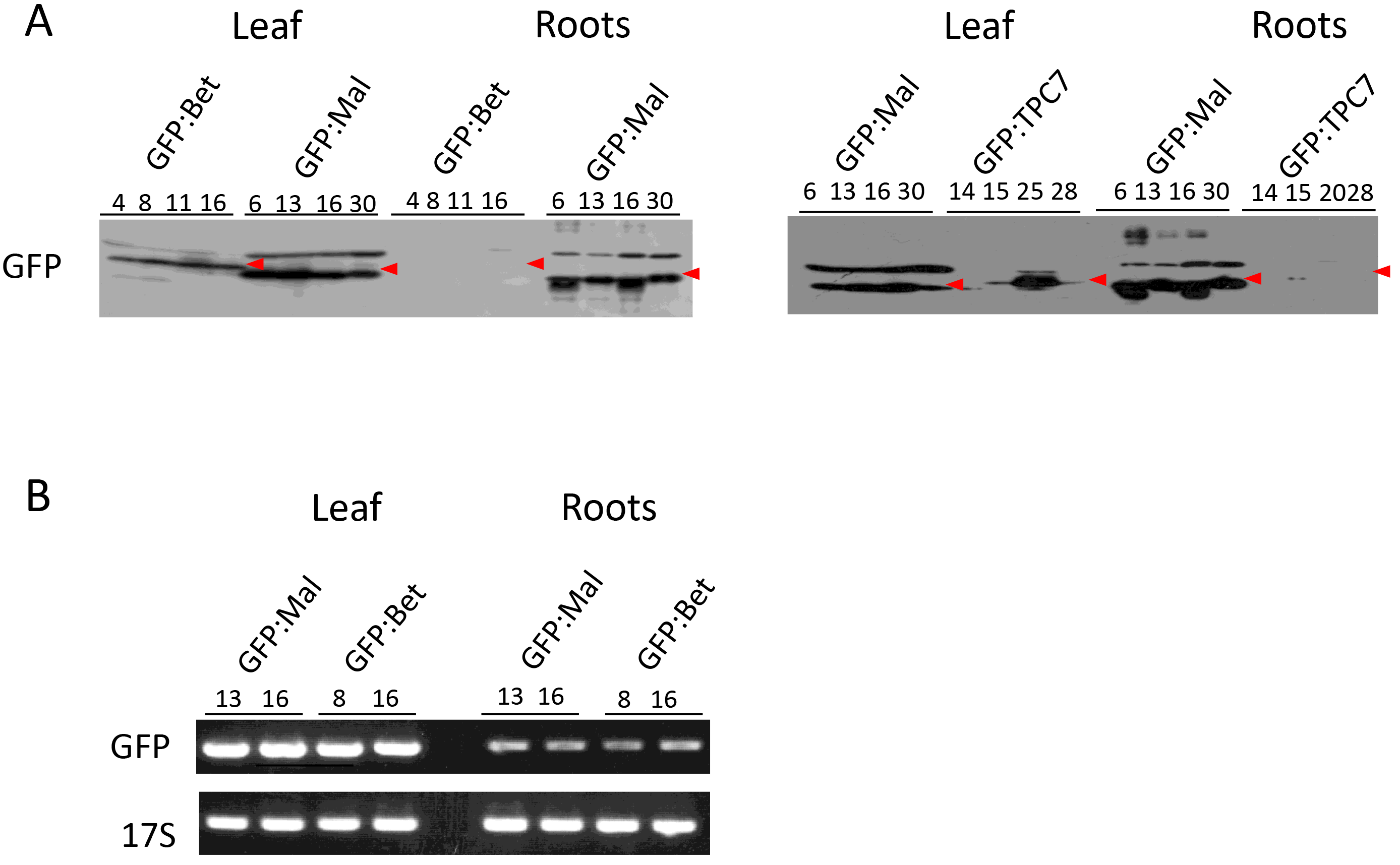
Expression of GFP:Mald1, GFP:Betv1 and GFP:TPC7 in leaf and roots of transgenic rice lines. **(A)** Immunoblot analysis of GFP:Mald1, GFP:Betv1 and GFP:TPC7 expressed in leaf and roots of transgenic rice lines under the control of the maize ubiquitin promoter. Processed 37-38 kDa fragments are indicated by arrow head. **(B)** RT-PCR of GFP:Mald1 and GFP:Betv1 transcripts in leaf and roots. Total RNAs were extracted from leaf and roots. The 17S rRNAs were used for normalization of starting RNA concentration in RT-PCR analysis.

When the transcript (mRNA) levels in leaf or roots between the Mal d 1 and Bet v 1 were examined by RT-PCR, there was little difference in their transcript levels between them, indicating that their accumulation levels are determined by post-transcriptional level (Fig. 4B).

### Subcellular localization of GFP:Mald1

When the GFP:Mald1 was expressed under the control of the maize ubiquitin promoter, it was detected in the whole grain under blue UV fluorescence (Fig. 2C). When this transgenic rice seed was vertically sectioned and observed using the confocal laser scanning microscopy, high fluorescence signal of GFP was detected in aleurone layer and embryo (Fig. 2D). Then, the subcellular localization of GFP:Mald1 in various parts of transgenic rice seed was examined in detail by fluorescence of GFP reporter (Fig. 2E-H) or immuno-histochemical analysis by anti-GFP antibody (Fig. S3) using the confocal laser scanning microscope. In the aleurone cells, the GFP:Mald1 was shown to be predominantly localized in huge spherical protein bodies (PBs) with a dimeter of >5 μm (Fig. 2E-F), whereas in the starchy endosperm it was localized in PB-Is containing prolamins and small granules with a size of `≦`1 μm (Fig. 2F, Fig. S3L). In the embryo of maturing seed, it was predominantly localized in ER lumen (Fig. 2H, Fig. S3H,K). To investigate the participation of chaperon proteins in huge PB formation, intracellular localization of several chaperon proteins was analyzed. As shown in Fig. S3 (A,B,D,I), BIP1, BiP4/5, PDIL1-1 and PDIL2-3 were co-localized in huge peripheral PBs in the aleurone cells, whereas they are also localized in ER lumen and PB-Is in the starchy endosperm. When antibodies against CNX (Fig. S3F), OsTiP3 (Fig. S3G) and Cys-rich 14kDa prolamin (RM1) (Fig.3SL) were used as markers for subcellular compartments of ER, protein storage vacuole (PB-II) and ER-derived PBs (PB-Is), respectively, they were not observed to be over-lapped with huge PBs. These results indicate that huge PBs accumulating GFP:Mald1 are formed by aid of BiPs and PDILs chaperone proteins.

Next, when intracellular localization of the GFP:Mald1 was examined in root, it was shown to be mainly localized as small granules in ER lumen (Fig. S4), since CNX, PDIL1-1, BIP1 were co-localized with the GFP:Mald1 at the peripheral sites of individual vegetative cell. Huge PBs could not be observed in root tissues. Strong fluorescence of GFP was observed at rooting site.

### Identification of regions of Mal d 1 responsible for stable accumulation in vegetative tissues

To investigate which domain(s) in the Mal d 1 molecule is involved in high and stable accumulation in vegetative tissues, the Mal d 1 was progressively deleted from the N or C termini and their resultant regions were fused to the GFP reporter as shown in Fig. 5A. They were expressed under the control of the maize ubiquitin constitutive promoter.

**Figure 5.**
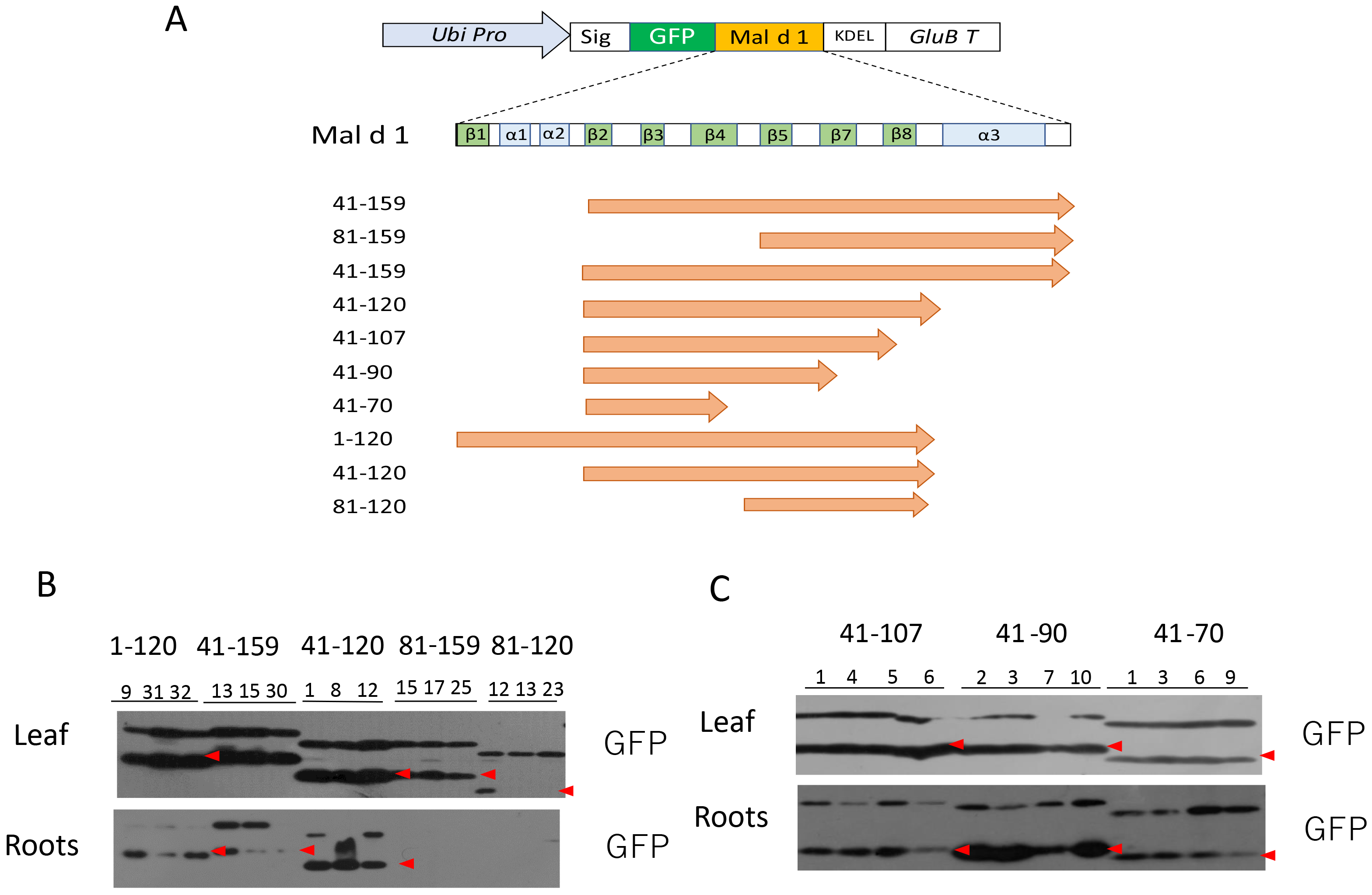
Deletion analysis of Mal d 1 molecule. **(A)** Schematic representation of the gene constructs used for deletion analysis of Mal d 1. α, α helical structure; β,β sheet structure **(B-C)** Effect of deleted Mal d 1 on accumulation levels of linked GFP in leaf and roots of transgenic rice. Total proteins extracted from leaves and roots of 4-5 representative independent transgenic lines for each expression construct were subjected to immunoblot analysis using anti-GFP antibody.

When these fusion product levels were examined in vegetative tissues (leaf and roots) by immunoblotting using anti-GFP antibody, the regions of the Mal d 1 between positions 1-120, positions 41-159 and positions 41-120 conferred higher level of accumulation than the regions between positions 81-120 and positions 81-159 (Fig. 5B). This result indicates that the region between positions 41-120 is critical to confer high level of accumulation in vegetative tissues. Then, to characterize the minimum region required for high expression in vegetative tissues, accumulation levels from the regions between positions 41-107, 41-90 and 41-70 were examined. As shown in Fig 5C, the region between positions 41-90 led to the highest accumulation level. Notably, further deletion of 20 amino acids (region between positions 41-70) or addition of 17 amino acids (region between positions 41-107) resulted in reduction of accumulation levels. These results indicate that the minimum region implicated in high accumulation in vegetative tissue is localized between positions 41-90. This region contains ‘P-loop’ (phosphate-binding loop) motif, (GxGGxGxxK) rich in glycine residue between positions 48-54 that is conserved in most of PR10 proteins (Fig. 1). In order to examine its function, this sequence (GNGGPGTIK) was substituted to ANAAPATIK. However, mutation of this sequence did not have any effect on accumulation levels of the linked GFP in most of leaves as well as seeds (Fig. S5). This result indicated that P-loop motif is not responsible for enhancing the accumulation level of the Mal d 1 in transgenic rice.

Next, it was examined how above deletions of the Mal d 1 have an effect on accumulation levels of the linked GFP in transgenic rice seeds (Fig. S6). Progressive deletion from the N terminus reduced the linked GFP accumulation levels (Fig. S6A). The region between positions 41-90 gave rise to higher level accumulation than the regions between positions 41-159, positions 41-120 and positions 41-70 (Fig. S6B). Furthermore, the region between positions 41-120 led to higher levels of accumulation than the regions between positions 81-120 and positions 1-120 (Fig. S6C). These results are fundamentally similar to those observed in vegetative tissues. Taken together, the region between positions 41-90 is involved in stable and high accumulation in seed as well as vegetative tissues.

### Domain swapping between the Mal d 1 and Bet v 1

To identify the domain that is implicated in high accumulation in vegetative tissues of transgenic rice harboring GFP:Mald1, domain swapping was carried out between the Mal d 1 and Bet v 1 as shown in Fig. 6A. When the C terminal regions of the Bet v 1 between positions 90-159 and positions 127-159 were exchanged with the corresponding regions of the Mal d 1, accumulation levels of these chimeric products (Bet:Mal90, Bet:Mal127) were very low in leaves like the native Bet v 1. That is, accumulation levels of these chimeric products were not significantly altered by fusion with the C terminal region of Mal d, indicating that the C terminal half region of the Mal d 1 has little enhancing effect. On the other hand, when the N terminal half region of Bet v 1 between positions 1and 90 was exchanged with the corresponding region of the Mal d 1, much higher level of this chimeric product (Mal:Bet90) than the native Bet v 1 was observed in leaves. These findings indicated that the N terminal half region of the Mal d 1 is responsible for high accumulation of the Mal d 1 in vegetative tissues. Therefore, in order to ascertain that the region between positions 41-90 is critical for high accumulation in vegetative tissue, the corresponding region was mutually exchanged between the Mal d 1 and Bet v 1. The Bet:Mal:Bet chimeric product, in which the region between positions 41-90 was substituted with the corresponding region of the Mal d 1, gave rise to high levels of accumulation in leaves of most of transgenic lines, whereas the Mal:Bet:Mal chimeric product containing the Bet v 1 sequence between positions 41-90 resulted in lower levels of accumulation in most of transgenic rice leaves. These results indicated that the region between positions 41-90 is mainly responsible for high accumulation in vegetative tissues of Mal d 1.

**Figure 6.**
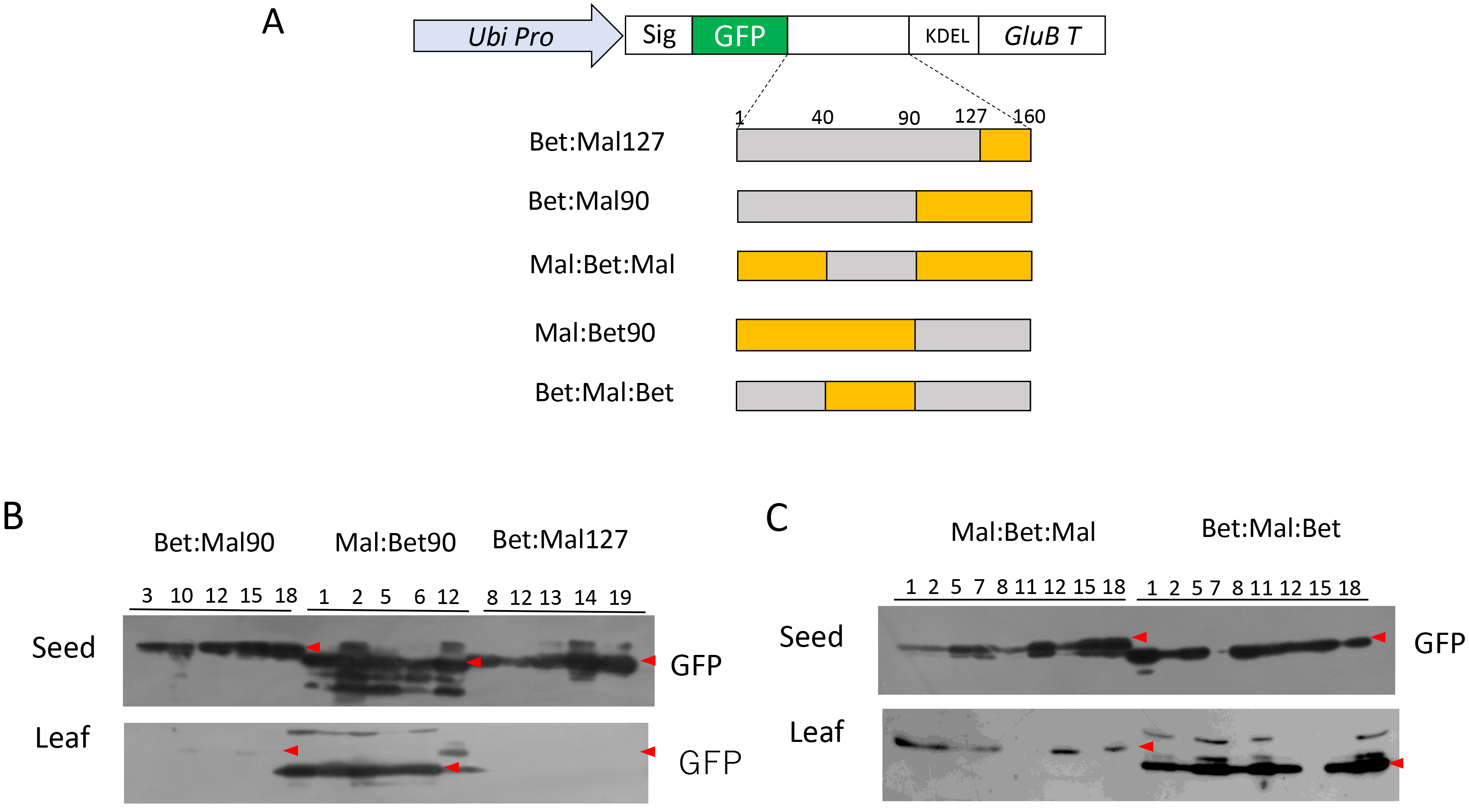
Domain swapping between the Mal d 1 and Bet v 1. **(A)** Schematic representation of gene constructs used for domain swapping. **(B-C)** Immunoblot analysis of various fusion products extracted from mature seeds and leaves of transgenic rice plants harboring various expression constructs using anti-GFP antibody.

### Aggregation property of Mal d 1

When the Mal d 1 containing KDEL ER retention tag was expressed under the control of the endosperm-specific glutelin *GluB-1* promoter, it was shown to be accumulated as an oligomeric form in a similar manner to the TPC7 and Bet v 1 that are deposited into huge PBs in endosperm (Fig. 7).

**Figure 7.**
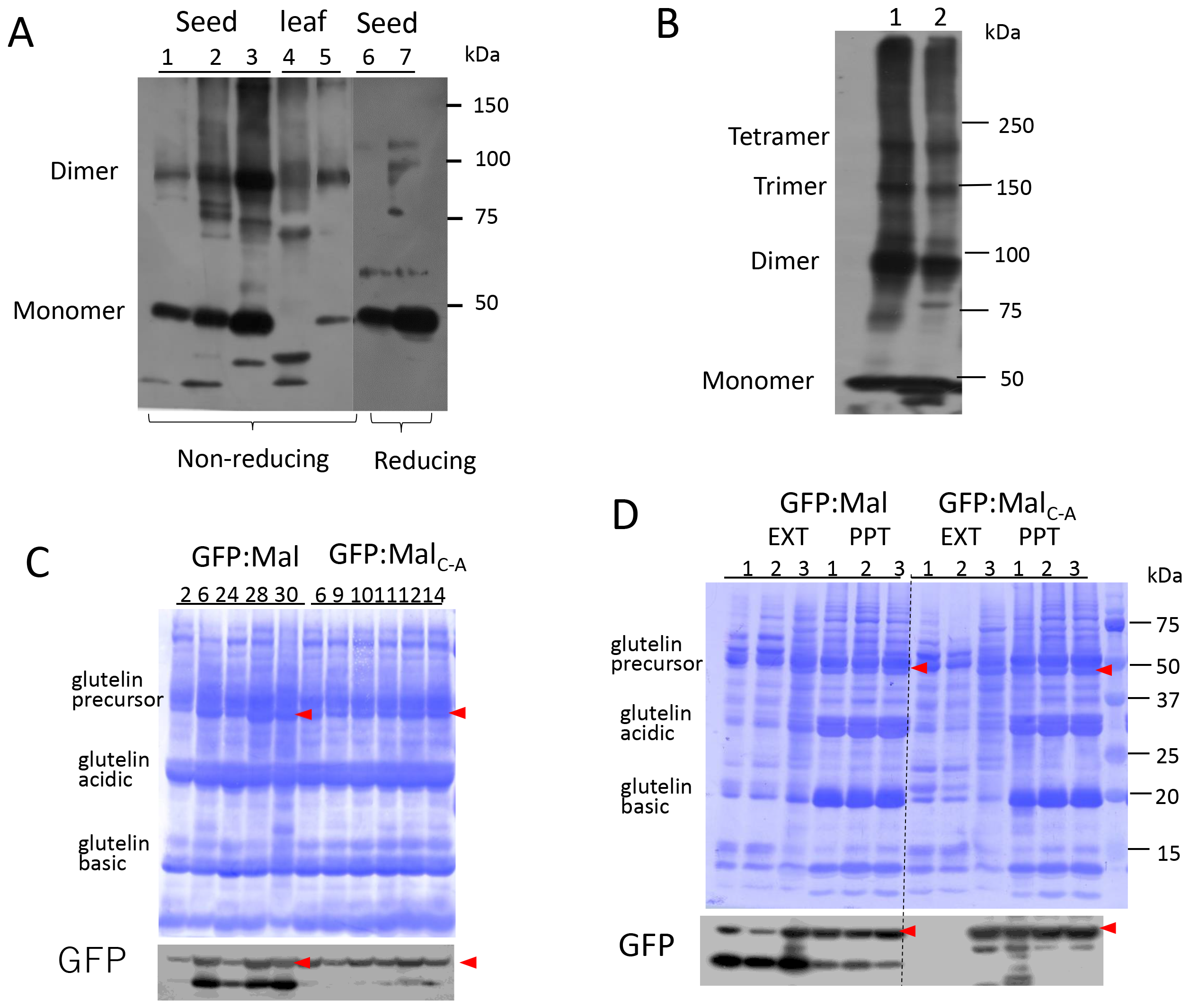
Comparison of aggregation property between GFP:Mald1 and GFP:Mald1_C-A_. **(A)** Immunoblot analysis of GFP:Mald1 and GFP:Mald1_C-A_ expressed in transgenic rice seeds and leaves. They were extracted with 50mM Tris-HCl (pH7.5) and 1%SDS without reducing agent and then separated by 8% SDS-PAGE. 1. Seed extract from transgenic rice harboring GluP::GFP:Mal 2. Seed extract from transgenic rice harboring UbiP::GFP/Ma_IC-A_ 3. Leaf extract from transgenic rice harboring UbiP::GFP:Mal, 4. Leaf extract from transgenic rice harboring UbiP::GFP:Mald1_C-A_. 5 and 6. Seed extracts of GluP::GFP:Mal and UbiP::GFP:Mal were treated with reducing agent before loading to SDS-PAGE. Aggregates were detected by immunoblotting using anti-GFP antibody **(B)** Separation of seed extracts of transgenic rice harboring GluP::GFP:Mal (1) and UbiP::GFP:Mal (2) by 6% SDS-PAGE. **(C)** Comparison of accumulation levels in transgenic rice seeds between GFP:Mald1 and GFP:Mald1_C-A_. **(D)** Difference in extraction efficiency from mature seeds and leaves of transgenic rice harboring UbiP::GFP:Mal and UbiP::GFP:Mal_C-A_ that were treated with 0.2%SDS (1), 0.5%SDS (2) and 1%SDS (3) in 50mM Tris-HCl (pH7.5). EXT, extracted; PPT, not extracted Arrow head represents the expressed GFP:Mald1 and GFP:Mald1_C-A_.

Next, to examine whether there is a difference in oligomeric formation among various tissues, the oligomeric formation of the GFP:Mald1 directed by the glutelin endosperm-specific or ubiquitin constitutive promoter was compared by separation on 8% non-reduced native PAGE. As shown in Fig.7A, dimer of GFP:Mald1 fusion protein was observed in the extracts from these transgenic rice seed and leaf. Furthermore, higher aggregates such as trimer and tetramer were also observed by separation on 6% non-reducing native PAGE (Fig. 7B). Such aggregates are suggested to be formed by self-aggregation via disulfide bond, because there is only one free cysteine (Cys) residue in the Mad d 1 like the Bet v 1 and TPC7. Therefore, the Cys residue in the Mal d 1 was substituted to the Ala residue to eliminate the Cys residue, which was then produced as a fusion protein with GFP in transgenic rice. When this GFP:Mal_C-A_ was extracted from seed and leaf and then analyzed by native PAGE, dimeric form of the modified Mal d 1 was observed irrespective of the absence of Cys residue (Fig. 7A line2 and 5). However, the migration of this aggregate was slightly different from that of the inherent one.

To examine whether this difference is attributed to the aggregate form, these GFP fusion proteins were extracted from transgenic seeds with different concentration of SDS (0.2%, 0.5% and 1%(w/v)). The GFP:Mald1 could be extracted with all the concentrations of SDS used here, whereas the GFP:Madl_C-A_ could not be extracted with 0.2% or 0.5% SDS buffer (Fig. 7D). This finding suggested that Cys free modified GFP:Mald1 is aggregated in different manner from the native one.

### Extraction of GFP:Mald1 from transgenic rice plants

GFP:Mald1 was extracted from nature seed and leaf of transgenic rice plants with various extraction buffers. Different concentration of SDS (0.2% and 1%) and 1% Tween20 and 1%Triton-X100 were used for solubilizing membrane proteins. When expressed in the endosperm under the control of the glutelin promoter, higher concentration of SDS (1%) was required for its extraction from mature seed. However, this extraction efficiency was much lower than the SDS-urea buffer. On the other hand, when expressed under the control of the ubiquitin promoter, it could be easily extracted from seed and leaf with all types of buffer used here. It is notable that extraction efficiency was almost same to the SDS-urea buffer. These results may be related to that the GFP:Mald1 directed by the maize ubiquitin promoter is localized in aleurone layer of seed and in ER lumen of leaf.

Next, to examine the cause of low extraction efficiency of GFP:Mald1 from transgenic rice seed that was directed by the endosperm-specific glutelin promoter, mature seed was treated with various extraction buffers containing reducing agent (10mM DTT) and various detergents. As shown in Fig. 8B, extraction efficiency of the GFP:Mald1 was improved by the presence of reducing agent (DTT). Furthermore, there is little difference in extraction efficacy by the presence or absence of KDEL ER retention sequence. These results suggest that the Mal d 1 interacts with Cys-rich prolamins via disulfide bond in starchy endosperm.

**Figure 8.**
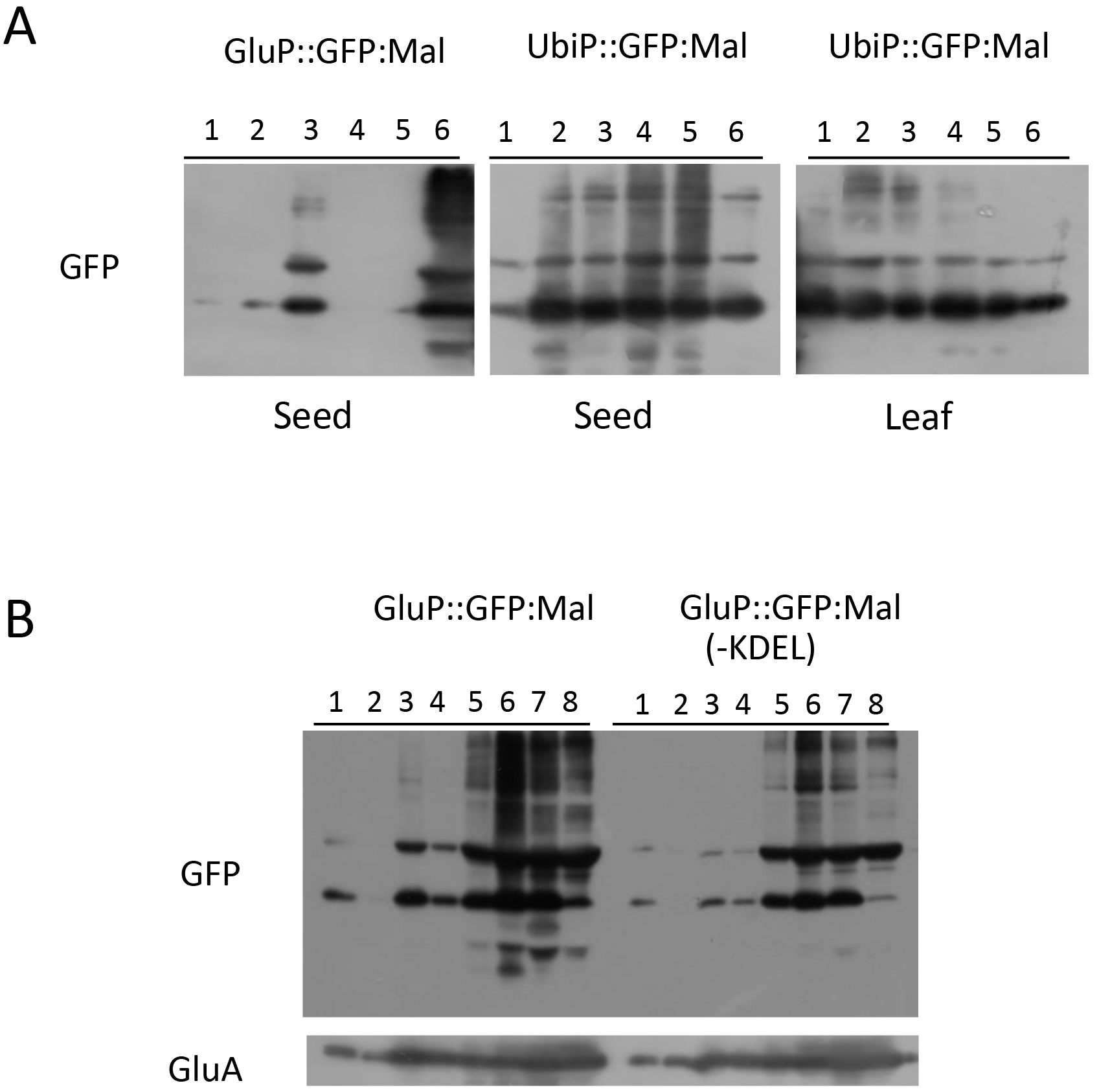
Extraction of GFP:Mald1 from seed and leaves of transgenic rice. **(A)** Seed powder and frozen leaf powder of transgenic rice harboring GluP::GFP:Mal or UbiP::GFP:Mal construct were treated with saline buffer (1) and saline buffer containing 0.2% SDS (2) 1% SDS (3) 1% TritonX-100 (4) and 1%Tween-20 (5) and urea-SDS buffer (6) **(B)** Seed powder of transgenic rice harboring GluP::GFP:Mal or GluP::GFP:Mal-KDEL) was treated with saline solution containing 1%CTAB (1), 10mM DTT (2), 1% SDS (3), 0.2% SDS and 10mM DTT (4), 0.5% SDS and 10mM DTT (5), 1% SDS and 10mM DTT (6), 6M urea and 10mM DTT (7) and urea-SDS buffer (8). Extraction efficiency by these treatments was examined by immunoblotting using anti-GFP and anti-GluA antibodies.

## Discussion

TPC7 and TPC 9 are derivatives of major birch pollen allergen Bet v 1, which have been developed through *in vitro* random recombination by means of DNA shuffling as versatile recombinant hypoallergenic allergen against multiple Fagales pollen allergens (Wallner *et al.* 2007). Since the TPC7 and TPC9 exhibit lower allergenicity and higher immunogenicity than the native Bet v 1, they are expected to act as the ideal tolerogens for allergen-specific immunotherapy against the major pollen allergens of trees belonging to the Fagales order Bet v 1 family. Therefore, to create rice seed-based oral allergy vaccine against the pollen allergy of Fagales order, we have generated transgenic rice accumulating the Bet v 1 and its derivatives, TPC7 and TCP9. We previously reported that high amounts of TPC7 (540 ug/seed) and native Bet v 1 were deposited in spherical huge protein bodies in endosperm cells of transgenic rice seeds, when expressed as secretory protein under the control of glutelin endosperm-specific promoter (Wang *et al.* 2013; Ogo *et al.* 2014).

It has been reported that more than 70% of birch pollen allergy patients develop an IgE-mediated hypersensitivity reaction termed as OAS after apple consumption (Geroldinger-Simic *et al.*, 2011). This is mainly attributed to the immunological cross-reaction between the Bet v 1 specific IgE and apple major allergen Mal d 1, because the Bet v 1 and Mal d 1 share high homology in the primary sequence. Pollen allergen Bet v 1 is specifically expressed in pollen, while food allergen Mal d 1 is expressed in vegetative tissues. Therefore, it is important to examine how these allergens are deposited in various tissues of transgenic rice, when these allergens are produced in transgenic rice and then used as rice-based allergy vaccines against these pollen and food allergy diseases.

As shown in Fig. 3, when the GFP:Betv1, GFP:TPC7 and GFP:Mald1 were expressed as secretory protein by ligating the N terminal signal peptide and C terminal KDEL tag under the control of the maize ubiquitin promoter, accumulation levels of GFP:Betv1 and GFP:TPC7 were very low in various vegetative tissues such as leaf, root and seed embryo except for seed (endosperm). In contrast, the GFP:Mald1 highly accumulated not only in seed, but also in leaf and root. When intracellular localization of the GFP:Mald1 was examined in maturing transgenic seeds, it is mainly deposited in huge PBs in aleurone cells and in ER-derived PBs (PB-Is) containing prolamin storage proteins in starchy endosperm cells. This accumulation pattern was remarkably different from that directed by rice endosperm-specific glutelin promoter, in which the GFP:Mald1 is predominantly deposited in huge PBs in the starchy endosperm cells. On the other hand, huge PBs were not observed in vegetative tissues, although high amounts of GFP:Mald1 accumulate in vegetative tissues. Then, taking these results into consideration, we examined why the GFP:Mald1 can be deposited into huge bodies in aleurone cells, but not in endosperm and vegetative tissues under the control of the maize ubiquitin promoter.

It has been suggested that huge PB formation is associated with the aggregate property of the recombinant proteins (Ogo *et al.*, 2014). Furthermore, expression levels may be also important for the formation of huge PBs, since maize ubiquitin promoter directs stronger expression in aleurone layer than in starchy endosperm of maturing seed, when examined by expression of the GUS reporter gene driven by the same maize ubiquitin promoter (Takaiwa *et al.*, 2007). Requirement for high accumulation of recombinant proteins above a threshold level has been pointed out for the formation of PB in elastin-like polypeptide (ELPs) and hydrophobin fusion proteins (Gutierrez *et al.*, 2013).

PB formation is suggested to be efficiently initiated by ER retention via the C terminal KDEL signal of GFP:Mald1. Actually, localization of GFP:Mald1 in ER lumen is observed in various tissues including aleurone layer and seed embryo, since small granules with green fluorescence are observed along ER membrane network all over the whole cell, which are connected as ER network. That is, the GFP:Mald1 is synthesized on rough ER, and subsequently transferred into the ER lumen. The size of PB gradually increases through binding of small aggregates, resulting in generation of huge spherical PB with a maximum size of >10 um. As shown in Fig. 9, a single or a few huge spherical PBs are finally formed by gathering several small size GFP:Mald1 aggregates in one cell. Aggregates are observed to be concentrated into a single huge PB through ER network connection. Finally, a single huge spherical PB is formed in many aleurone cells. Growth in PB size is accompanied by an increase in accumulation levels. Furthermore, PB formation is known to be mediated by aid of some chaperon proteins such as BiPs and PDILs that are implicated in folding and assembly. When subcellular localization of some BiPs and PDILs were examined by immuno-histochemical analysis, these chaperons are localized within huge PBs containing GFP:Mald1 as shown in Fig. S5, indicating that huge PBs in aleurone cells are formed in a similar manner as huge PBs observed in the starchy endosperm (Ogo *et al.* 2014). Aleurone cells are rich in protein storage vacuoles (PSVs) referred as aleurone bodies and lipid bodies. Some storage proteins such as rice embryo globulin-2 proteins (REG2) and phytic acid are accumulated inside aleurone bodies. Antibodies against OsTiP3 and CNX as PSV and ER marker are not localized to huge PBs in the aleurone cells. Huge PBs are suggested to be independently formed by self-aggregation of Mal d 1. Generation mechanism of this huge PB is fundamentally same to that of Zera, ELPs and hydrophobin-I in plant cells (Conley et al, 2011; Saberianfar and Menassa, 2017).

**Figure 9.**
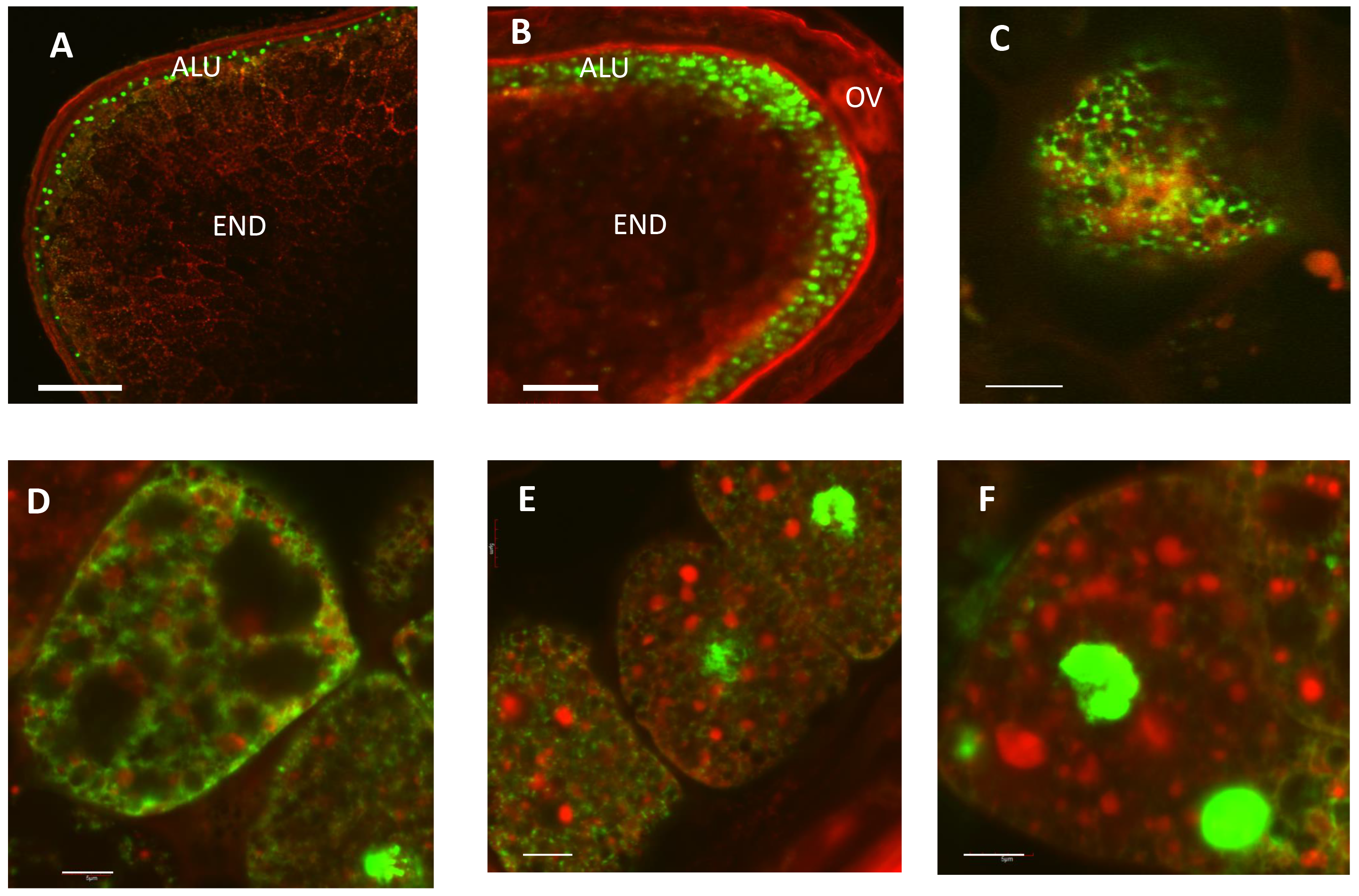
Subcellular localization of GFP:Mald1 in aleurone cells. Green fluorescence by GFP represents the localization of GFP/Mald1. Red represents the rhodamine B stained particles. **(A)** Ventral site of sectioned seed **(B)** Dorsal site of sectioned seed **(C-F)** Huge PB formation process of GFP:Mald1 in alurone layer. ALU, aleurone layer; END, endosperm; EMB, embryo; OV, ovular vascular trace. Thick bars= 200 μm Thin bars= 5 μm

Bet v 1 and Mal d 1 belong to the same PR10 group and share about 55-68% amino acid sequence homology to each other. However, it is notable that their accumulation levels in various tissues are quietly different even when expressed under the control of the same constitutive ubiquitin promoter. The GFP:Mald1 highly accumulates in various vegetative tissues and seed, whereas high accumulation of the GFP:Betv1 is limited to the endosperm and its accumulation level in the vegetative tissues is relatively low. It is notable that transcript levels between the GFP:Mald1 and GFP:Betv1 in leaves and roots are almost same, when examined by RT-PCR using GFP primers. Post-translational regulation is involved in such difference in accumulation levels. Non-correlationship between transcript levels and accumulation products have been reported for several seed storage proteins in maize aleurone cells (Reyes *et al.*, 2011). By contrast, foreign recombinant protein such as 7Crp peptide selectively accumulated in endosperm rather than other tissues (embryo and vegetative tissues) when expressed under the constitutive promoter (Takaiwa *et al.*, 2007). Difference in intracellular localization or post-translational modification such as glycosylation may be also implicated in such post-translational regulation.

In this study, we first demonstrated by deletion analysis of the Mal d 1 that the region between positions 41-90 is mainly responsible for high and stable accumulation in vegetative tissues. Participation of this region in high accumulation in vegetative tissues was further confirmed by domain swapping between the Mal d 1 and Bet v 1. As shown in Fig. 6, the exchange of this Mal d 1 region between positions 41-90 with the corresponding region of Bet v 1 (Bet:Mal:Bet) resulted in higher accumulation of the linked GFP in vegetative tissue than the native Bet v 1. This region includes very conserved P element motif and N-glycosylation site of Bet v 1 and TPC7. Moreover, we previously reported by domain swapping between TPC7 and Bet v 1 that the sequence between positions 32-160 is important for the formation of huge and high number of PBs (Ogo *et al.*, 2014). This result also supports the contribution of C terminal half region to the huge PB formation. Furthermore, this region has been suggested to be involved in immunological cross-reactivity between Mal d 1 and Bet v 1, since this region was identified as conformational B-cell epitope (discontinuous epitopes) by antibody binding (Mirza et al., 2000). This epitope is formed by the segment between Glu42 and Thr52 along with Arg70, Asp72, His76, Ile86 and Lyn97 of Bet v 1. The tertiary structure of this region exhibits antiparallel (3-sheet structure (β-2, (β-3, (β-4 and (β-5), which covers a large proportion of the inner cavity surface (Ahammer *et al.*, 2017). Mutagenesis of conserved amino acid within this region (E45S) also reduced IgE binding (Spangfort *et al.*, 2003). These evidences also suggest that the region between positions 41-90 is structurally important and is localized on the protein surface of the Mal d 1..

When the GFP:Mald1 was specifically expressed under the control of the endosperm-specific promoter, it was difficult to extract it from mature seed. This is due to that it is deposited into huge ER-derived PBs by self-aggregation or PB-Is by interaction with Cys-rich prolamins via disulfide bonds (Takaiwa *et al*, 2009). Higher concentration of detergent (SDS) and reducing agent (DTT) are required for efficient extraction of Mal d 1 from transgenic rice seed. By contrast, when expressed under the control of the constitutive promoter, the GFP:Mald1 can be easily extracted from various tissues. Especially, it can be extracted from transgenic mature seeds even with saline solution. This can be explained by the observation that the GFP:Mald1 is localized in ER lumen of vegetative tissue or accumulated mainly in huge PBs of aleurone cells.

Bet v 1 has been reported to generally form a mixture of monomers, dimers and higher order oligomers (Scholl *et al.*, 2005). Dimerization was also reported for Mal d 1 (Ma *et al.*, 2006). When GFP:Mald1 was produced in transgenic rice, it displays multimeric formation in seed and vegetative tissues (Fig. 8). The Mal d 1 has one cysteine residue like the Bet v 1. This cysteine residue is suggested to be implicated in disulfide bond formation for self-aggregation or interaction with the Cys-rich prolamins in transgenic rice seeds. However, even though this cysteine residue was substituted to Ala by mutagenesis, oligomeric (dimeric) formation was retained. Notably, this mutagenesis resulted in the alternation in physicochemical property of the accumulated modified Mal d 1, since it became difficult to extract the mutagenized Mal d 1 from vegetative tissues. The mutagenized Mal d 1 lacking Cys residue suggested to be falsely folded, leading to change in physicochemical property of Mal d 1. Therefore, cysteine residue in Mal d 1 molecules may be critical for the correct folding or oligomeric formation via disulfide bond.

It has been demonstrated in this study that localization and generation of such huge PBs in transgenic rice are dependent on the tissue specificity and strength of the promoter used for expression. When directed by the maize ubiquitin constitutive promoter, huge PBs are observed to be formed mainly in aleurone cells, whereas expression by the endosperm-specific glutelin promoter gives rise to their production in starchy endosperm. Higher expression above a threshold level is essentially required for huge PB formation. However, although many various foreign recombinant proteins have been produced in our laboratory, such huge PBs could not be formed except for the PR10 proteins such as Bet v 1 and Mal d 1 even when the same strong endosperm-specific promoter has been utilized, suggesting that the specific physicochemical property such aggregation property may be associated with the huge PB formation. Their tertiary structure leading to high aggregates may be responsible for the huge PB formation by helping to increase the accumulation levels. When the GFP was produced by fusion with Mal d 1, it was highly and stably accumulated not only in seed, but also in leaf and root (Fig. S7). Notably, accumulation level of GFP has been significantly increased by fusion with Mal d 1. Furthermore, GFP:Mald1 fusion protein can be more easily extracted from various tissues of transgenic rice than that specifically expressed in transgenic rice seeds. Therefore, Mal d 1 may be utilized as fusion partner for the production of high-value recombinant proteins in plant like Zera and ELPs. Further works will be required whether production of foreign recombinant proteins can be enhanced by fusion with Mal d 1.

## Acknowledgments

We thank Ms. K. Miyashita, Y. Ikemoto and Y. Yajima for technical assistance and Dr. Kenjirou Ozawa for encourage of this research.

## Supplementary Figures

**Figure S1. Immunoblot analysis of GFP:Mald1 fusion proteins expressed in transgenic rice seeds under the control of the endosperm-specific glutelin 2.3 kb *GluB-1* promoter.** Total proteins were extracted from mature seeds with SDS-urea buffer, and then separated by 13% SDS-PAGE. Proteins were visualized with CBB staining and accumulation level of GFP:Mald1 was analyzed by immunoblotting using anti-GFP or anti-TPC7 antibody. Effect on expression of chaperon proteins by accumulation of GFP:Mald1 were examined by immonoblotting using several anti-chaperone proteins.

**Figure S2.Immunoblot analysis and SDS-PAGE of GFP containing KDEL tag expressed as secretory protein in transgenic rice seed under the control of the ubiquitin promoter.** GFP accumulates in both embryo and endosperm of mature seed.

**Figure S3.Subcellular localization of GFP:Mald1 in aleurone and starchy endosperm cells in maturing seeds at 15-18 DAFs.** Green signal indicates GFP fluorescence or immunofluorescence of GFP immune-stained with anti-GFP antibody. Red signal indicates immunofluorescences of OSBiP-1, OSBiP4/5, OSPDIL1-1, OSPDIL2-3, CNX, OsTiP3 and RM1 (Cys-rich 14kDa prolamin) immunostained with their specific anti-rabbit antibodies. Signals were detected using second antibody with the Alexa-488-conjugated goat anti-mouse IgG (green) and Alexa-564-conjuated goat anti-rabbit IgG (red). The images correspond to the merge channels resulting from the combination of anti-mouse IgG recognizing GFP (green) and anti-rabbit IgG recognizing chaperons (red). Co-localization is represented by yellow or orange color. Scale bars are 5 μm. ALU, alurone layer; END, endosperm; EMB, embryo

**Figure S4.Intracellular localization of GFP/Mald1 and various chaperone proteins in root tissue.** Transverse section of transgenic rice root. The images correspond to the merged channel resulting from the combination of GFP of GFP:Mald1 (shown in green) and chaperon proteins (shown in red). Colocalization is shown in orange. Bars 5 μm

**Figure S5.Effect of mutation of Mal d 1 P-loop motif on accumulation level in leaf and seed.** P-loop motif of Mal d 1 was mutagenized and this modified Mal d 1 was expressed as fusion protein with GFP in leaves and seeds of several independent transgenic rice lines (GFP:Mal(P)). Accumulation levels in these leaves and mature seeds were compared with those of GFP:Mald1 as control.

**Figure S6. Effect of deleted Mal d 1 on accumulation levels of linked GFP in seeds of transgenic rice.** Total proteins extracted from mature seeds of 4-5 representative independent transgenic lines for each expression construct were subjected to immunoblot analysis using anti-GFP antibody

**Figure S7. Comparison of accumulation levels of GFP, GFP:Betv1 and GFP:Mald1 in leaf and seed.** GFP, GFP:Betv1 and GFP:Mald1 were expressed under the control of the maize ubiquitin promoter.

## Abbreviations

Cys: cysteine
BiP: binding protein
ER: endoplasmic reticulum
ER: elastin-like polypeptide
GFP: green fluorescence protein
OSA: oral allergy syndrome
PB: protein body
SDS-PAGE: sodium dodecyl sulfate polyacrylamide gel electrophoresis
RT-PCR: reverse transcription-polymerase chain reactions
PR10: pathogen related protein 10
PDIL: protein disulfide isomerase

